# Optimized Lipid Extraction and Hybrid Annotation Pipeline for Individual Chitinous Mesozooplankton Using UPLC-HRMS

**DOI:** 10.1101/2025.09.24.678324

**Authors:** Jiwoon Hwang, Bethanie R. Edwards

## Abstract

High-resolution lipidomics at the scale of individual mesozooplankton offers a powerful tool for understanding trophic interactions and carbon cycling in marine ecosystems, but chitinous exoskeletons present challenges for efficient lipid extraction. Here, we develop and validate an optimized Bligh and Dyer–based extraction protocol that incorporates in-line glass bead homogenization, yielding a 2.5-fold increase in lipid recovery. When combined with an increased injection volume, this protocol achieves a 4.4-fold gain in signal intensity. This workflow enables the robust detection of intact lipid species from single *Calanus* copepods, eliminating the need for additional homogenization equipment or extended extraction steps, thereby making it broadly accessible for analytical applications.

To address the limitations of current annotation pipelines, we directly compared adduct-hierarchy (LOBSTAHS) and fragmentation-based (MS-DIAL) approaches, finding systematic biases that reshape lipidomic profiles depending on the computational strategy employed. Additionally, by integrating a wax ester-specific fragmentation library, we demonstrate improved annotation of marine-relevant lipid classes largely absent from conventional databases. Together, this extraction and hybrid annotation pipeline enables high-resolution, compound-specific lipidomics of individual mesozooplankton, capturing biological heterogeneity while remaining scalable to pooled samples. Our approach provides a critical methodological advance for tracing lipid metabolism across trophic levels and for quantifying the role of mesozooplankton lipids in marine biogeochemical cycles.

## Introduction

The advent of high-resolution mass spectrometry coupled with ultra-high performance liquid chromatography (UPLC-HRMS) has enabled the detection and quantification of increasingly more compounds in less biomass, with single-cell lipidomics beginning to emerge as a tractable technique (Hunter et al., 2021; Wang et al., 2023). However, the smallest biological unit for which reliable and routine lipidomic measurements have been reported is the individual multicellular eukaryotic organism, like those living at the micro- and mesoscale. In the marine ecosystem, small invertebrates at this level often occupy basal positions in food webs and serve as key intermediaries in the transfer of essential lipids (i.e., lipids that are not produced by most metazoans and must be obtained from their diet) - such as docosahexaenoic acid (DHA) and eicosapentaenoic acid (EPA) - to higher trophic levels, underscoring the importance of lipidomic profiling at this scale.

### Lipid extraction challenges in chitin-encased organisms

Many of these organisms, including copepods and other small crustaceans, possess chitinous exoskeletons. Chitin, a nitrogen-containing polysaccharide, has a supramolecular structure, stabilized by extensive inter- and intramolecular hydrogen bonding that renders it insoluble in common organic solvents (Rinaudo, 2006). Because chitin is the most abundant amino polysaccharide in nature, this insolubility presents a significant challenge to researchers looking to extract the lipidome of chitin-encased species. Currently, most extraction methods for such taxa rely on heavily extended versions of traditional lipid extraction protocols that employ longer processing steps such as shaking, sonication, or incubation. These methods, while effective, are often time-consuming, labor-intensive, depend on large quantities of biomass, require extra equipment, or are otherwise not optimized for organisms encased in chitin (Supp. Table 1, Figure 1).

**Figure 1.**
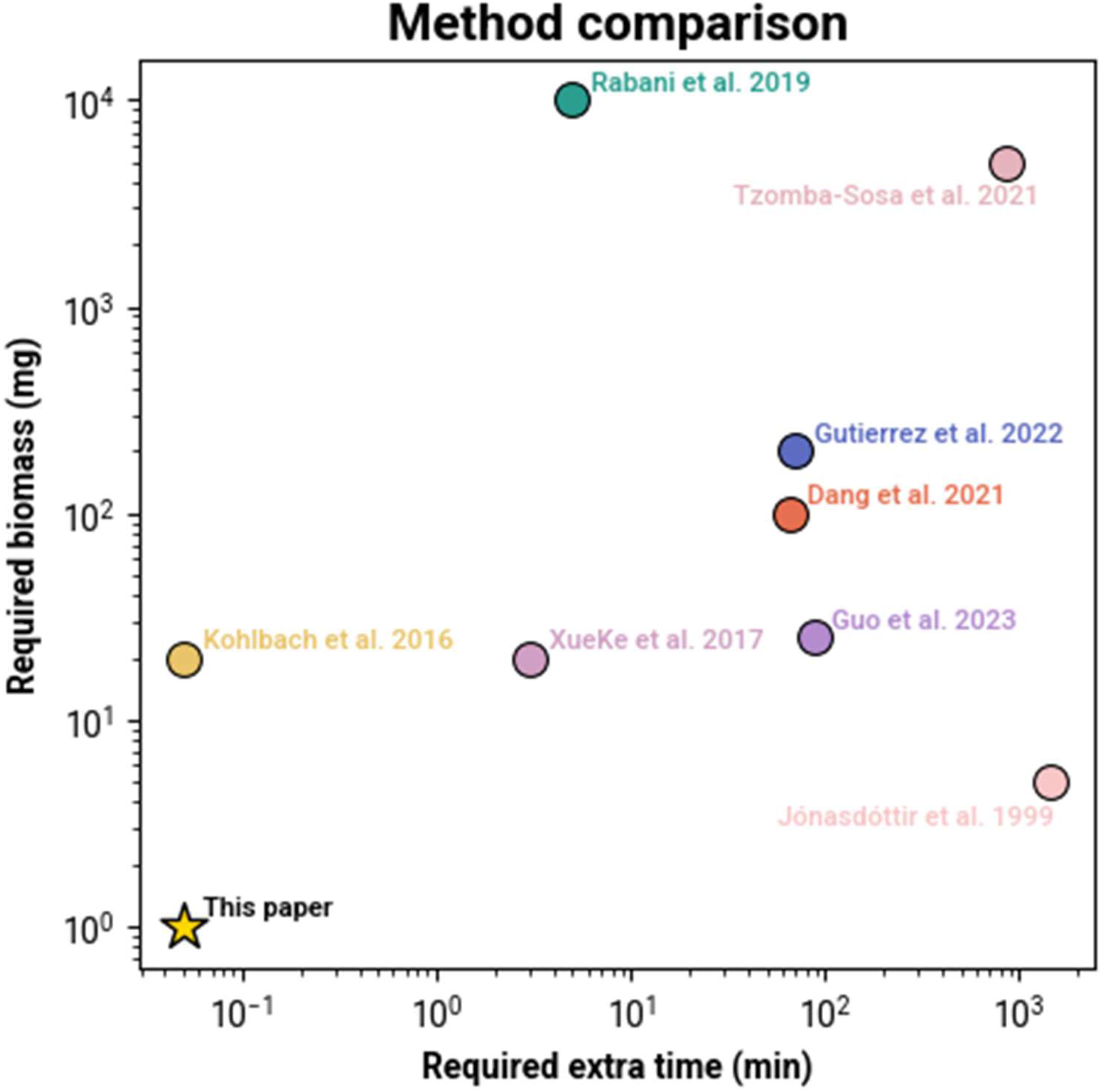
Comparison of lipid extraction methods applied to various chitin-bound organisms. Both axes are presented on a log₁₀ scale. Only data points for which both time and corresponding biomass measurements were available are included.

**Table 1.** Overview of treatments and sample groups.

| Number of<br>individuals<br>Treatment | n=1 | n=4 | n=6 |
| --- | --- | --- | --- |
| Glass beads | Yes | No | 607<br>Yes608 |
| Increased injection<br>volume | Yes | No | 609<br>No 610<br>611 |
| Control (+optimized<br>LC method) | Yes | Yes | 612<br>Yes613<br>614 |

The original Bligh and Dyer method was first developed in the 1950s to extract lipids from fish muscle. In this system, the lower organic phase contains lipids and other relatively non-polar compounds, while the upper aqueous phase retains more polar metabolites. The protocol was modified in the 2000s by substituting chloroform for dichloromethane (DCM) to reduce toxicity and improve handling (Cequier-Sánchez et al., 2008; Sturt et al., 2004; Van Mooy et al., 2008) and was subsequently optimized for the extraction of marine phytoplankton (Popendorf et al., 2013; Van Mooy and Fredricks, 2010). Here, we present a hybridized lipid extraction method adapted from Van Mooy and Fredericks’ version of the classic Bligh and Dyer protocol, that enhances extraction efficiency from *Calanus* spp., a genus of cosmopolitan zooplankton belonging to the class Copepoda. Other versions of the Bligh and Dyer method have been applied to analyze copepod metabolomes (Hansen et al., 2013; Mayor et al., 2015), as well as one instance of a Matyash lipid extraction, that uses Methyl tert-butyl ether (MTBE) instead of chloroform or DCM, on commercially sourced *Calanus* copepod specimens (Matyash et al., 2008; Wood et al., 2023). Given the physiological diversity within copepods and the differences in experimental context, direct comparisons with our results are limited. For these reasons, we focus on the development and optimization of the Bligh and Dyer protocol adapted for marine microalgae, as established and applied within our research group.

Our method, inspired by bead-beating techniques commonly used in DNA extraction protocols, reduces overall processing time compared to existing protocols and can be integrated directly into the pre-existing lipid extraction workflow without the need for additional extraction steps. While ceramic beads are commonly used in molecular biology, we use glass beads for their proven compatibility with lipidomics work due to their non-reactivity with organics solvents and combustibility. We demonstrate a 2.5-fold increase in average extraction efficiency, and when paired with our analytic optimization (i.e., the increased injection volume), a 4.4-fold improvement in average signal intensity (peak area) per unit concentration.

### Current approaches to lipid identification in marine systems

While UPLC-HRMS offers high resolution and the ability to detect a broad range of spectral features within a sample, accurate compound annotation remains a challenge. In algal and marine lipidomics, identification of compounds typically relies on a multi-pronged strategy: (1) MS/MS fragmentation patterns, and (2) the “adduct hierarchy” of lipids, which infers compound identity and deconvolutes isomeric and isobaric compounds based on preferred ionization behavior during electrospray ionization (ESI), and (3) comparison of retention times to authentic standards. Although all approaches are widely used and experimentally valid, each presents specific limitations.

For example, adduct-based identification tools, such as LOBSTAHS (Lipid and Oxylipin Biomarker Screening Through Adduct Hierarchy Sequences), are constrained by empirical validation of adduct hierarchies for additional compound classes, and regioisomers within the database resulting in competing annotations. In contrast, MS/MS-based annotation methods such as MS-DIAL or MZmine, especially those employing *in silico* libraries, can overcome this limitation but require substantial amounts of data generation. Fragmentation spectra can also vary significantly depending on collision energy, as well as other instrument-specific parameters (Lee et al., 2024). As databases expand, the computational cost and time for annotation increase exponentially, further limiting accessibility to researchers without access to high-performance computing resources.

Furthermore, most publicly available lipid databases are built around mammalian, terrestrial or freshwater lipidomes and lack coverage of marine-specific lipid classes. For instance, our study focuses on wax esters, a group of relatively neutral lipids that are produced by the ester bonding of a fatty acid and an alcohol. While wax esters are one of the predominant lipid classes in *Calanus* copepods, making up to 90% of the total lipid mass (Albers et al., 1996), they are, to our knowledge, notably absent from commonly used databases. We therefore highlight the usage of an “18-series” fragmentation pattern that has been observed in wax esters and β-hydroxy ceramides (Chen et al., 2015; Tsugawa et al., 2017), but has not been implemented in pre-existing lipid databases (Kind et al., 2013; Tsugawa et al., 2020).

To illustrate the practical consequences of these limitations, we conducted a direct comparison between two widely used annotation platforms: LOBSTAHS and MS-DIAL. This comparison highlights how differences in database structure, spectral libraries, and computational pipelines can significantly influence annotation outcomes, even when applied to identical datasets. Our results reveal significant divergence between the tools, necessitating a hybridized approach that combines elements from both platforms. However, reconciling output from disparate peak-picking algorithms and annotation strategies adds to the complexity and time required for data processing. While the widely used GNPS (Global Networks Public Social Molecular Networking), especially when paired with MZmine, exemplifies the value of community-curated spectral libraries and network-based annotation (Schmid et al., 2023; Wang et al., 2016), it is mainly focused on terrestrial and mammalian compounds, limiting our ability to comprehensively annotate untargeted marine lipidomes. We advocate for a unified, community-driven effort to develop a comprehensive, marine-specific spectral database, analogous to the centralized reference systems used in genomics (e.g., INSDC (International Nucleotide Sequence Database Collaboration)).

## Materials and Methods

### Sample collection

Copepods from the genus *Calanus* were collected from Svalbard, Norway aboard the *R/V G.O. Sars* using a 180 µm mesh size Bongo net during the 2022 Nansen Legacy Polar cod cruise in November, with a smaller subset collected in February 2023. Samples were all identified under an on-deck stereomicroscope and flash frozen in centrifuge tubes at -80 ℃ using liquid nitrogen. The second subset was collected on a privately chartered boat with a WP2 net, from 180 m to the surface. Collected copepods were kept at -2 ℃ and brought back to the University Centre in Svalbard (UNIS), after which they were transported by plane to Oslo at 0 ℃. This second subset of samples were part of a larger experiment that involved three weeks of lab culture at a pH of 7.9 (control conditions) before being flash frozen for analysis at the University of California, Berkeley.

### Lipid Extraction & Analysis

The experimental design is summarized in Table 1 and Figure 2. In brief, “control” groups containing either 1, 4, or 6 individuals underwent a Bligh and Dyer extraction, followed by injection of 2 µL of the extract into the UHPLC for analysis. Two types of “treatment” conditions were tested. Treatment A involved mechanical disruption using a bead-beating protocol and was applied to both individual samples (n = 1) and pooled samples (n = 6). Treatment B involved increasing the UHPLC injection volume to 10 µL and was applied only to individual samples.

**Figure 2.**
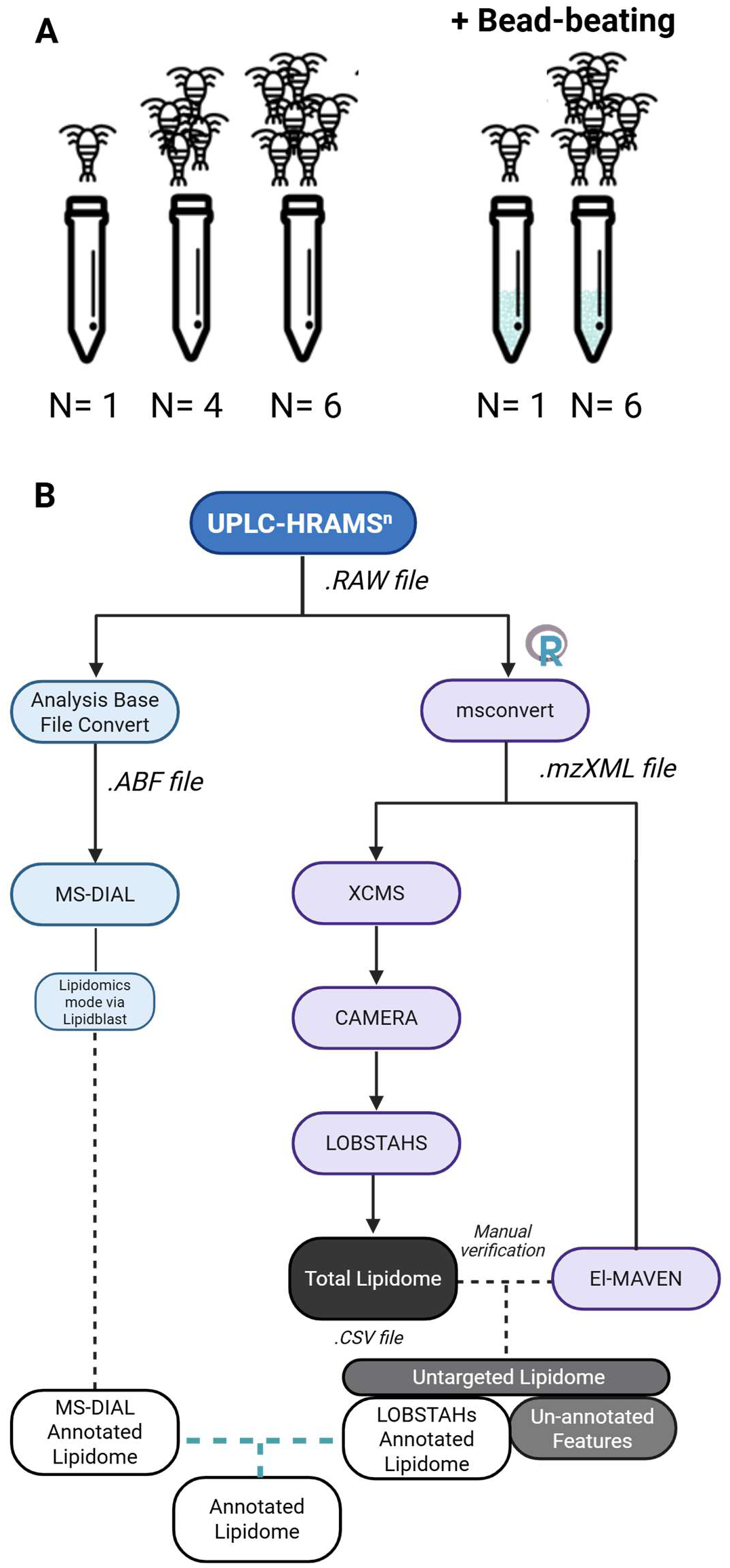
(A) Depiction of sample groups. Numbers of individuals in each group are denoted underneath. **(B) Lipid annotation pipeline comparing MS-DIAL and LOBSTAHS.**

Lipids were extracted using a modified Bligh and Dyer protocol adapted from Popendorf et al (2011). Individual, partially thawed copepods were transferred from cryovials to the inner walls of pre-cleaned glass centrifuge tubes using a clean metal spatula. For Treatment A, approximately 2 mL of combusted glass beads (0.5mm diameter, QIAGEN, Germantown, MD, USA) were added to each tube. A solvent mixture of methanol, dichloromethane (DCM), and phosphate-buffered saline (PBS) in a 2:1:0.8 (v/v/v) ratio was added, along with 10 µL of internal standard (EquiSplash LIPIDOMIX® Quantitative Mass Spec Internal Standard (Avanti Polar Lipids, Alabaster, AL, USA) diluted 2:1 from stock in methanol). Tubes were placed in an ultrasonic water bath for 10 minutes. Subsequently, additional DCM and PBS (1:1, v/v) were added to induce phase separation, followed by centrifugation at 1600 rpm for 10 minutes at 4 °C. The lower organic phase was collected, evaporated under a stream of nitrogen gas, and reconstituted in 70:30 acetonitrile:isopropanol (ACN:IPA, v/v). Extracts were stored at -80 °C until UHPLC-HRMS analysis.

The typical sample injection volume for reverse-phase UHPLC-MS/MS on a Vanquish UHPLC with an Accucore C8 column (155 mm x 2.1 mm x 2.6 µm) in tandem with an Orbitrap ID-X mass spectrometer (all from Thermo Scientific, San Jose, CA, USA) was 2uL. For Treatment B, the analytical run was performed with a 5-fold increase in injection volume (10 µL). We used a chromatographic method modified from Hummel et al. (Hummel et al., 2011) to include an additional isocratic hold, optimized for wax ester separation based on preliminary analyses using wax ester standards (Figure 3, Supplementary Table 2). Briefly, eluent A was liquid chromatography-grade water with ammonium acetate and acetic acid as additives, and eluent B was 70:30 ACN: IPA with the same additives. The starting gradient was 45/55 (A/B), which held for 1 min before shifting to 1/99 over 17 minutes, followed by an isocratic hold at 1/99 for 13 minutes. During the first minute of the mass spectrometric method, early-eluting small compounds not retained on the column (also colloquially known as the solvent front) were diverted to waste to remove compounds too hydrophilic to be retained that elute at the start of the run. Compounds were then analyzed in positive mode at a resolution of 120,000. MS/MS was analyzed as data-dependent acquisition for compounds greater than 1E+04 signal intensity. A mass inclusion list was used to acquire MS3 of compounds losing a free fatty acid fragment at the time that triacylglycerols (TAGs) and wax esters elute (Kiyonami et al., n.d.).

**Figure 3.**
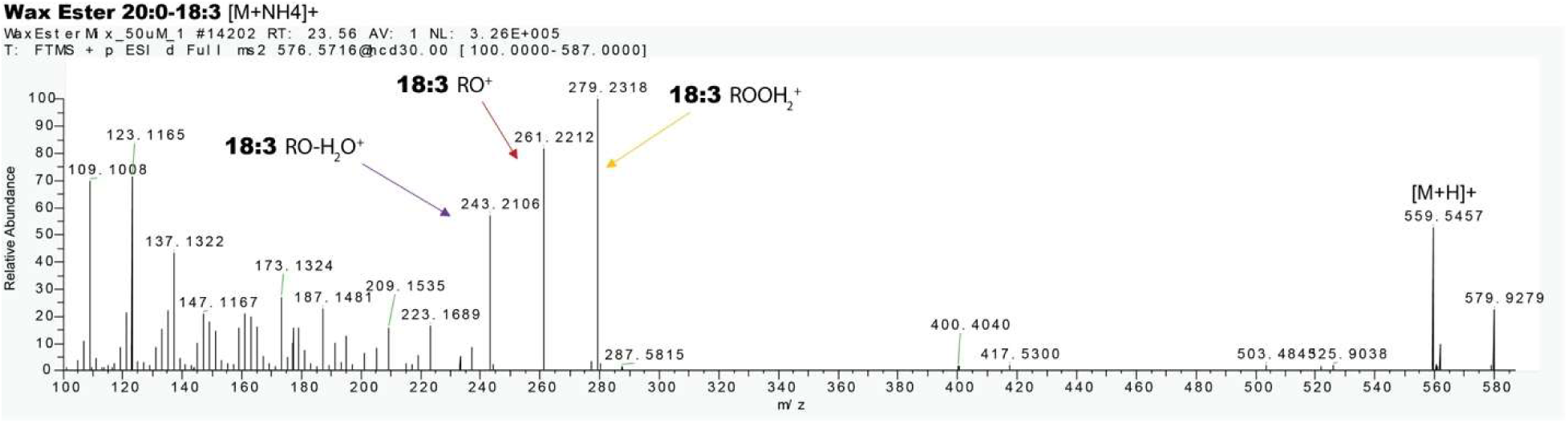
MS/MS fragmentation of an Arachidyl linolenate wax ester standard. Here, the dash-notation for wax esters, where the fatty alcohol R’ and fatty acid R are separated by a dash (R’-R), is used instead of the slash-notation, where R and R’ are denoted as R/R’.

Compounds from all samples were putatively annotated using the LOBSTAHS platform and default lipidomics database. For comparative analysis, the same samples were also processed using MS-DIAL using the default lipidomics database and identical peak detection parameters. Raw UHPLC-MS^n^ data was converted to mzXML files for LOBSTAHS annotation and abf files for MS-DIAL. Only peaks above a threshold of 5E+4 were picked in both analyses, with a 0.005ppm m/z threshold and a 0.2 min retention time threshold for peak grouping and library matching. The annotation workflow is detailed in the supplementary text of Hwang et al., 2025 (Hwang et al., 2025) and the computational load of workflows are compared in Supplementary Table 3.

Wax esters initially annotated in LOBSTAHS were further verified based on their characteristic “18-series” fragmentation pattern using MS-DIAL’s ability to query custom databases. This pattern involves the progressive loss of water molecules from the fatty acid moiety, transitioning from [RCOOH₂]⁺ to [RCO]⁺ and subsequently to [RCO-H₂O]⁺ (Figure 3; Chen et al., 2015). We created a custom fragmentation library (msp format) containing wax ester species composed of fatty acid and alcohol chains with the following parameters: (1) carbon chain lengths ranging from C₈ to C₃₀, (2) 0-8 double bonds, and (3) 0-4 oxygen modifications. In our assessment of wax ester annotations in our dataset, we focused on compounds containing 20:1 and 22:1 fatty alcohols, which are known to be synthesized *de novo* by polar copepods (Kattner and Hagen, 1995; Sargent et al., 1977). Additionally, a 36:5 wax ester was included because its unusual composition (8:0/28:5) nevertheless produced a clear 18-series fragmentation pattern. This made it a useful benchmark compound for testing the robustness of our custom fragmentation library across both typical and atypical wax ester species. The reduction of detected features to annotated and high-confidence compounds is summarized in Table 2 and Supplementary Figure 1.

**Table 2.**
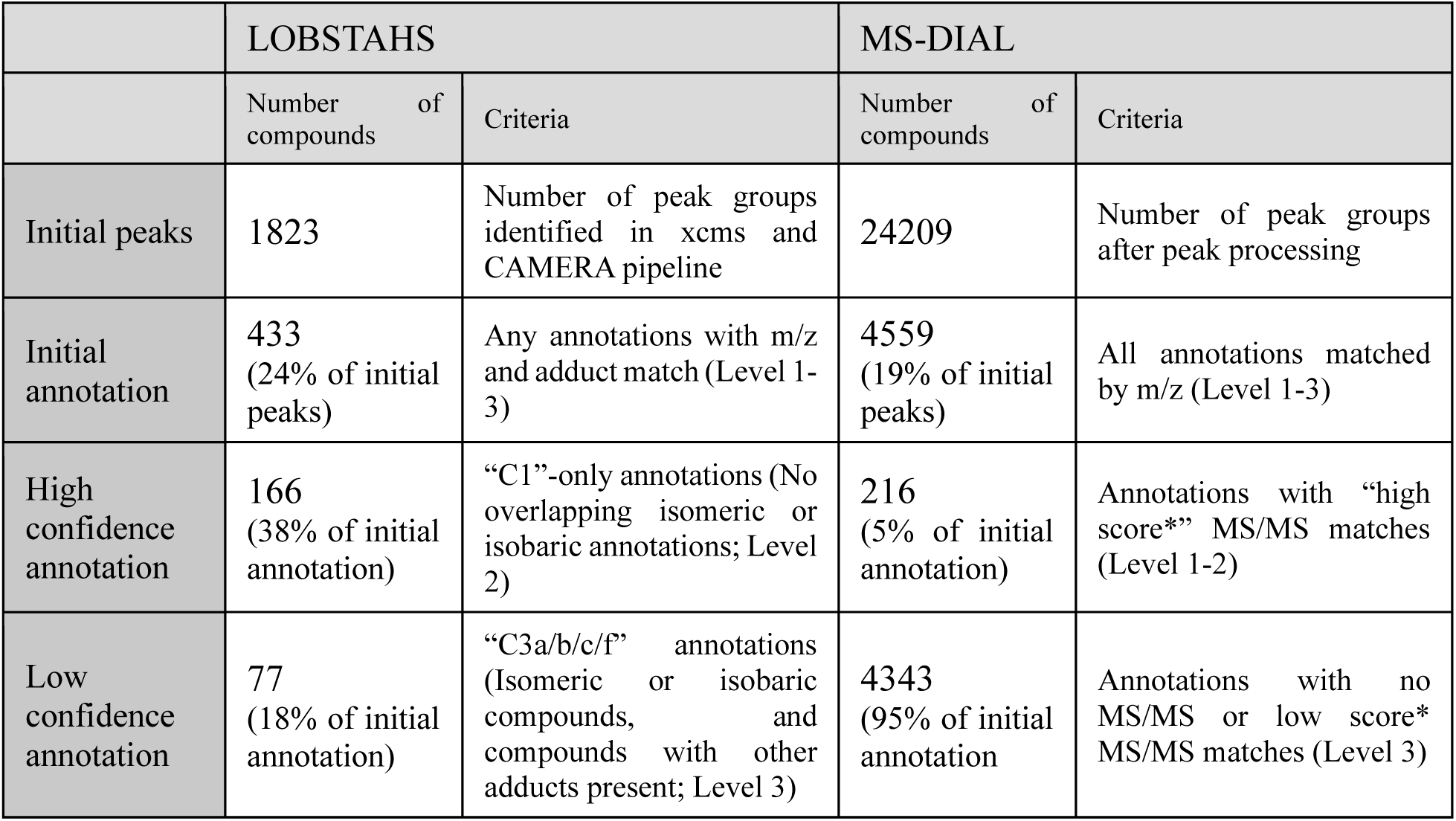
Data reduction table for LOBSTAHS and MS-DIAL annotation pipelines. Annotation “levels” correspond to annotation confidence levels put forth by the Chemical Analysis Working Group (CAWG) Metabolomics Standards Initiative (MSI); Level 1 is has a confirmed structure with reference standard and MS/MS and retention time match, Level 2 has a probable structure with library match, Level 3 has a putative class or compound group and Level 4 is an unknown compound (Sumner et al., 2007)

### Statistics

All statistical analyses were performed in R v4.4.2 or Python 3.12.8. To assess treatment effects on individual wax ester compounds (Table 3), we used one-way analysis of variance (ANOVA) (aov() in R), followed by Tukey’s HSD post-hoc comparisons (TukeyHSD(), α = 0.05). Results are reported as mean peak area ± standard deviation (SD) and standard error (SE), and significance was accepted at *p* < 0.05. Assessment of treatment effects in overall wax ester abundance (Figure 4A) was conducted via hierarchical clustering using Euclidean distance and Ward’s D2 linkage (hclust(method = "ward.D2")), which minimizes within-cluster variance and is well suited for quantitative lipidomic data (Murtagh and Legendre, 2014).

**Figure 4.**
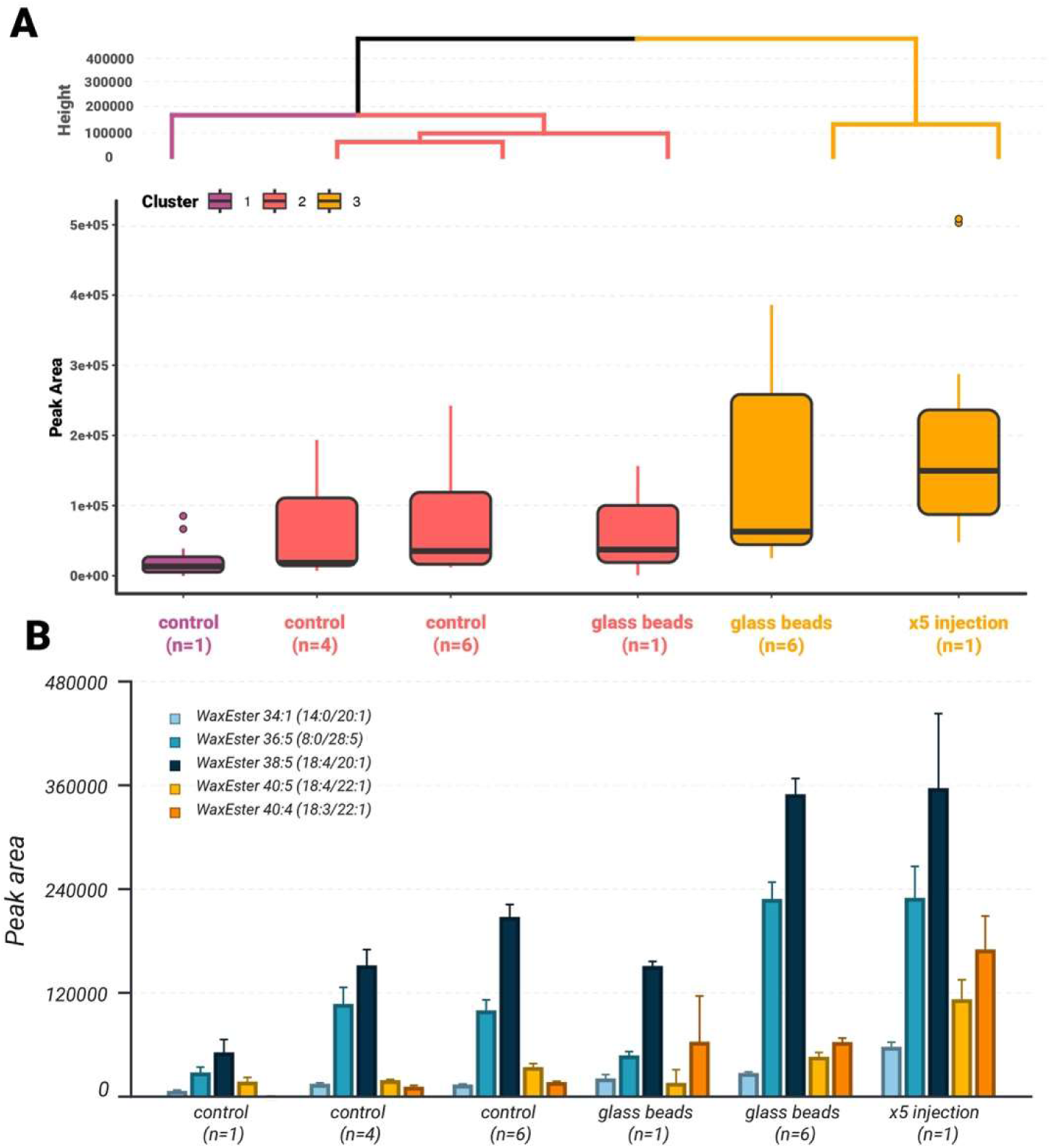
Relative abundance of wax esters under different treatment conditions. Treatment A is denoted as “glass beads” and treatment B is denoted as “x5 injection”. **(A) Average wax ester peak area across experimental treatments with hierarchical clustering.** Values represent the mean ± standard error of the mean (SEM) for each treatment (n as indicated). Treatments were clustered based on overall peak intensity using hierarchical clustering with Euclidean distance and Ward’s D2 linkage (Ward.D2 in R), which minimizes total within-cluster variance and is well suited for quantitative comparisons of peak-area data. **(B) Composition of individual wax ester species across treatment groups.** Each color represents a distinct wax ester, annotated by carbon:double bond count (e.g., 34:1) and its fatty acid/fatty alcohol composition (e.g., 14:0/20:1). Bars show the mean ± standard error of the mean (SEM) of peak area for each compound within each treatment.

**Table 3.** Overview of wax ester compounds of interest. Mean peak areas (± standard deviation and standard error) under different treatment conditions are shown. We tested for an overall effect of treatment on peak area with a one-way ANOVA in R (aov(), R 4.4.2), followed by Tukey’s HSD post-hoc comparisons (TukeyHSD(), α = 0.05). Significance was accepted at p < 0.05.

| compound | treatment | group | mean | sd | sem | n |
| --- | --- | --- | --- | --- | --- | --- |
| WaxEster 34:1 (14:0/20:1) | 1 2uL | a | 6633.705 | 2173.91597 | 1086.95798 | 4 |
| WaxEster 34:1 (14:0/20:1) | 4 2uL | a | 14744.9775 | 2449.56572 | 1224.78286 | 4 |
| WaxEster 34:1 (14:0/20:1) | 6 2uL | a | 13981.7 | 1607.30566 | 803.652829 | 4 |
| WaxEster 34:1 (14:0/20:1) | 1 glassbeads 2uL | ab | 21163.105 | 6119.46472 | 4327.115 | 2 |
| WaxEster 34:1 (14:0/20:1) | 6 glassbeads 2uL | b | 27287.1775 | 2785.82555 | 1392.91277 | 4 |
| WaxEster 34:1 (14:0/20:1) | 1 10uL | c | 57590.8775 | 10617.5092 | 5308.75459 | 4 |
| WaxEster 36:5 (8:0/28:5) | 1 2uL | a | 27956.755 | 12347.8746 | 6173.9373 | 4 |
| WaxEster 36:5 (8:0/28:5) | 4 2uL | a | 107177.36 | 37988.5308 | 18994.2654 | 4 |
| WaxEster 36:5 (8:0/28:5) | 6 2uL | a | 99663.5575 | 24427.3359 | 12213.6679 | 4 |
| WaxEster 36:5 (8:0/28:5) | 1 glassbeads 2uL | a | 47725.945 | 6122.08101 | 4328.965 | 2 |
| WaxEster 36:5 (8:0/28:5) | 6 glassbeads 2uL | b | 228615.105 | 38828.4858 | 19414.2429 | 4 |
| WaxEster 36:5 (8:0/28:5) | 1 10uL | b | 229848.875 | 72647.3987 | 36323.6994 | 4 |
| WaxEster 38:5 (18:4/20:1) | 1 2uL | a | 51199.8675 | 29687.1625 | 14843.5813 | 4 |
| WaxEster 38:5 (18:4/20:1) | 4 2uL | a | 151666.025 | 36765.0646 | 18382.5323 | 4 |
| WaxEster 38:5 (18:4/20:1) | 6 2uL | a | 207658.66 | 28911.7648 | 14455.8824 | 4 |
| WaxEster 38:5 (18:4/20:1) | 1 glassbeads 2uL | ab | 150964.7 | 7538.71995 | 5330.68 | 2 |
| WaxEster 38:5 (18:4/20:1) | 6 glassbeads 2uL | b | 349587.28 | 36255.3661 | 18127.683 | 4 |
| WaxEster 38:5 (18:4/20:1) | 1 10uL | b | 356703.108 | 172239.025 | 86119.5125 | 4 |
| WaxEster 40:5 (18:4/22:1) | 1 2uL | a | 17463.7625 | 9546.46914 | 4773.23457 | 4 |
| WaxEster 40:5 (18:4/22:1) | 4 2uL | a | 19001.6675 | 1689.87549 | 844.937745 | 4 |
| WaxEster 40:5 (18:4/22:1) | 6 2uL | a | 34259.1125 | 7841.04919 | 3920.52459 | 4 |
| WaxEster 40:5 (18:4/22:1) | 1 glassbeads 2uL | a | 15941.695 | 21440.9413 | 15161.035 | 2 |
| WaxEster 40:5 (18:4/22:1) | 6 glassbeads 2uL | a | 46225.6025 | 9632.03251 | 4816.01626 | 4 |
| WaxEster 40:5 (18:4/22:1) | 1 10uL | b | 112531.505 | 45405.3278 | 22702.6639 | 4 |
| WaxEster 40:4 (18:3/22:1) | 1 2uL | a | 372.59 | 745.18 | 372.59 | 4 |
| WaxEster 40:4 (18:3/22:1) | 4 2uL | a | 11426.7225 | 3099.69033 | 1549.84517 | 4 |
| WaxEster 40:4 (18:3/22:1) | 6 2uL | a | 16840.4975 | 1607.84469 | 803.922344 | 4 |
| WaxEster 40:4 (18:3/22:1) | 1 glassbeads 2uL | ab | 63429.2 | 74856.8836 | 52931.81 | 2 |
| WaxEster 40:4 (18:3/22:1) | 6 glassbeads 2uL | a | 63079.4325 | 9052.59849 | 4526.29924 | 4 |
| WaxEster 40:4 (18:3/22:1) | 1 10uL | b | 170119.443 | 77458.5607 | 38729.2804 | 4 |

To evaluate whether pooling samples altered lipidomic profiles, we performed partial least squares–discriminant analysis (PLS-DA) in Python using scikit-learn (PLSRegression) (Figure 5). Peak-area matrices were log₁₀-transformed (zeros replaced with the minimum finite value) and mean-centered by feature. Model performance was assessed by leave-one-out cross-validation (LOOCV) using a logistic classifier trained on the two-component scores. We report classification accuracy, sensitivity, and specificity from out-of-fold predictions. Global model fit was summarized by R²X (variance in *X* captured), R²Y (variance in *Y* explained), and Q² (LOOCV predictive ability). Statistical significance of group separation was evaluated with a 1,000-permutation test, where the permutation *p*-value was defined as the fraction of shuffled models with Q² greater than or equal to the observed Q².

**Figure 5.**
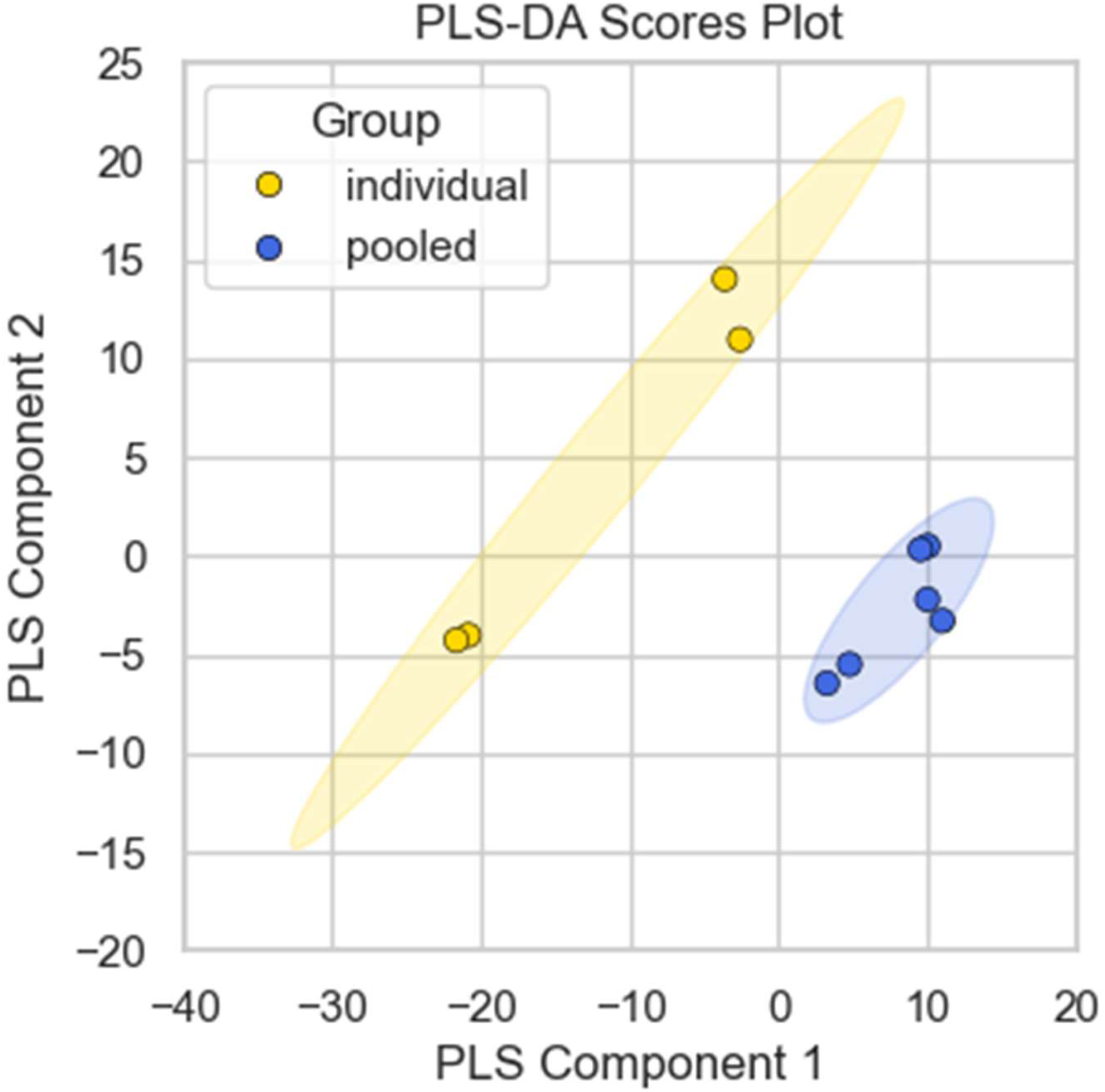
PLS-DA analysis of individual versus pooled samples. Two-component PLS-DA scores plot of log₁₀-transformed, mean-centered lipidomic profiles from individual (gold) and pooled (blue) samples. Each point represents one sample, and 95 % confidence ellipses are shown for each group. Component 1 captures 52.2 % and Component 2 captures 17.1 % of the X-variance (R²X = 0.522, 0.171). The model explains 98 % of the Y-variance (R²Y = 0.98) and achieves strong predictive performance (Q² = 0.93 by leave-one-out CV; permutation p = 0.007). Classification accuracy is 100 % (sensitivity = 100 %, specificity = 100 %).

## Results & Discussion

### Analysis of single copepods reveals robust recovery of lipids

In our control experiment, where non-bead-beat samples were run with 2 μL injection volumes, we found that, while single-copepod extractions still had detectable peak areas, many compounds were below the typical threshold that was used for peak detection in our peak-picking algorithms and programs (5E+4; Figure 4). We decided to increase the injection volume 5-fold, from 2 μL to 10 μL (Treatment B), instead of lowering the peak detection threshold. We deemed the second option non-viable due to issues such as lower-quality peaks, peaks with lower signal-to-noise ratios, and non-Gaussian peak shapes that ultimately complicate annotation further down the pipeline.

After assessment of these options, we opted to increase the injection volume. We were aware of potential ion suppression issues in higher injection volumes. However, we observed a non-linear increase in peak area, approximately 9-fold higher than the control value (Figure 4, Table 3). Under the same parameters, the peak-peaking algorithm XCMS discovered ∼8000 peaks in the 10 μL injection samples, compared to ∼4300 peaks found in the 2 μL injection samples, showcasing the effects of our signal amplification. Thus, we conclude that single copepod samples can be extracted to great efficiency.

The 4-individual and 6-individual groups produced large, well-resolved peaks, reflecting sufficient analyte concentration and eliminating the need for further method optimization. Notably, the 6-individual control group exhibited an average peak area only ∼3.6 times greater than that of the single-individual controls, a discrepancy primarily attributed to biological heterogeneity within the sample pool. This heterogeneity was evident in the PLS-DA analysis, which clearly showed the wider spread of biological variability within individual samples compared to pooled (Figure 5). These results indicate that individual biochemical profiles were well preserved in our analytical workflow, while pooled samples exhibited more cohesive, averaged trends.

### Glass beads greatly improve extraction efficiency

When comparing control and bead-beat (Treatment A) samples, pooled samples (n = 6) showed a 2-fold increase in peak area, while individual samples (n = 1) exhibited a greater effect, with a 2.9-fold increase (Figure 4). Extending sonication time would further improve yields for larger sample sets. Paired with the 1.8-fold increase from increased injection volume (Treatment B; volume-corrected), the combined effect of both treatments is estimated to yield a theoretical 3.6∼5.2-fold increase in peak area, excluding ion suppression effects. However, we emphasize that the objectives for each alteration differ; injection volume increase is a method to amplify the sensitivity or resolution of measurements (that should not affect post-analysis corrected peak areas to a large degree), while bead-beating results in an “actual” increase in absolute amount of extracted compounds.

As seen in Supplementary Table 1 and Figure 1, this is not the first attempt to extract lipids from chitinous organisms. However, our method has the smallest footprint in resources such as time, extra equipment (*i.e.*, cost efficiency and accessibility), and required sample size. It does not require any extra steps other than adding the combusted beads into centrifuge tubes before adding solvent. While homogenization using external equipment such as a Precellys24 bead homogenizer is effective, it requires quantitative transfer between two vessels and can therefore complicate full sample recovery. Other homogenizers that have been used for copepod analysis, such as Potter-Elvehjem homogenizers, are optimal for softer tissue samples and hence could require more initial biomass and/or longer agitation times to obtain an adequate concentration. Most other copepod lipid extraction methods are optimized for derivative analytical methods such as FAME (fatty acid methyl esterification), which differ in scope from our intact lipidome analysis.

### Annotation is strongly biased depending on the chemoinformatic pipeline utilized

Compounds from all control and treatment samples were comparatively identified using two different annotation pipelines; the adduct-hierarchical LOBSTAHS and fragmentation-based MS-DIAL, which have distinct default lipid databases (Supp. Figure 2). Overall, high confidence annotations within the copepod lipidome revealed a dominance of neutral lipids that mainly function as energy storage such as wax esters, and triacylglycerols (TAG), lipids that participate in metabolism such as coprostanol esters, and diacylglycerols (DAG), as well as phytoplankton-associated membrane lipids such as betaine-like lipids (BLL) and digalactosyldiacylglycerols (DGDG) (Figure 6). However, while the dominant lipid classes identified by each algorithm were broadly similar in functional category (energy storage as opposed to energy harvesting, structural, signaling), at the compound and lipid subclass level the annotated lipidomes varied considerably, resulting in distinct profiles that reflect differences in annotation method only. In the LOBSTAHS-annotated lipidome, wax esters and coprostanol esters were the most frequently detected lipid subclasses, whereas in MS-DIAL, TAGs and DAGs dominated the annotations (Figure 6A). Weighting the feature by peak area to reflect relative quantities of lipids did not alter the overall composition of the dominant lipid classes, but it did result in a modest reordering of their relative abundances (Figure 6B).

**Figure 6.**
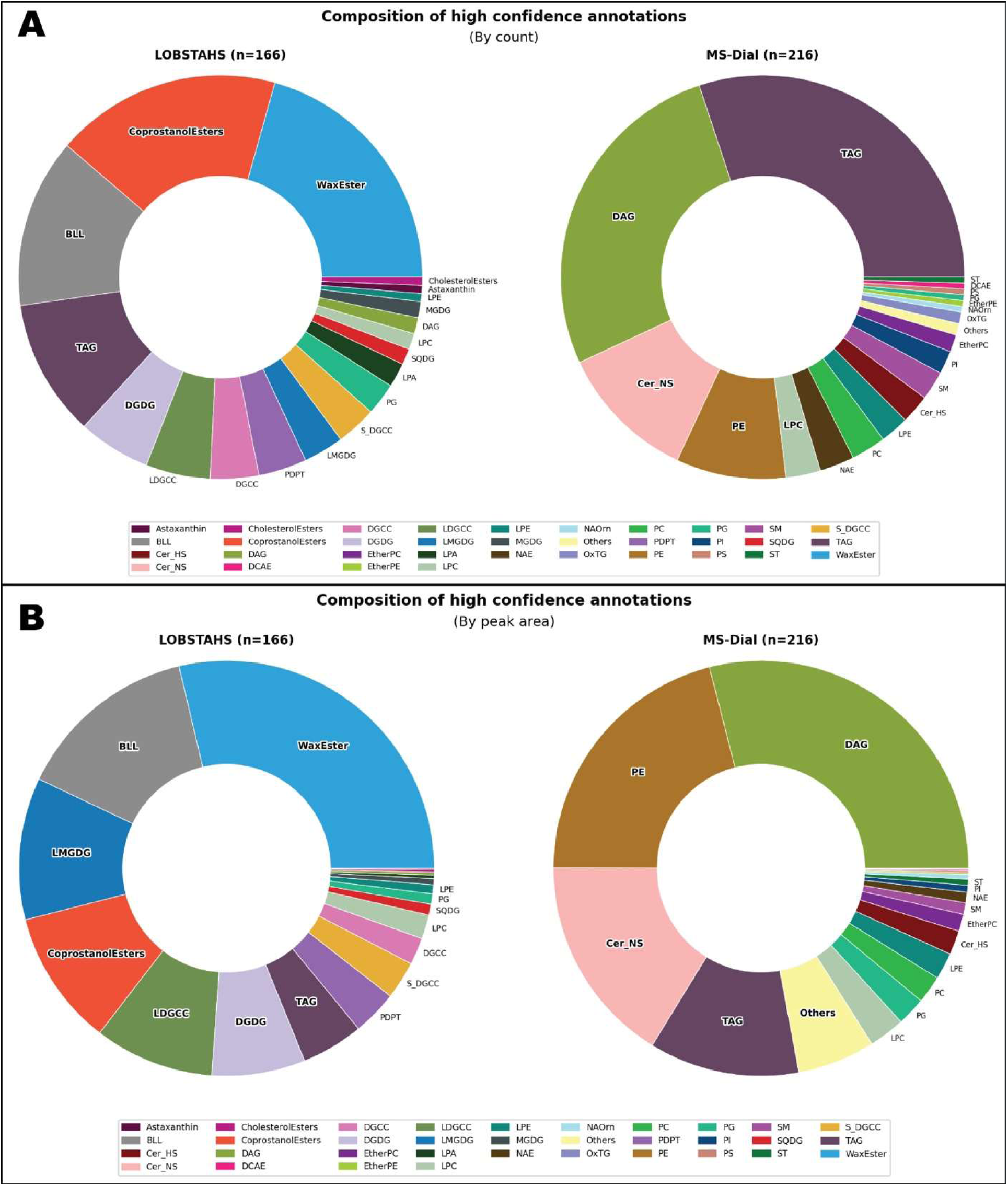
Lipid composition of high confidence (level 1-2) annotations in LOBSTAHS and MS-DIAL. All slices are ordered by abundance. (A) Size of slices are the number of compounds in each lipid group (B) Size of slices are the combined peak area of all compounds in each lipid group

It is worthwhile to note that DAGs, cholesterol esters, and wax esters are frequently co-annotated with ceramides and N-acyl ethanolamine (NAE) (Figure 7). Their simple elemental compositions, consisting primarily of carbon, hydrogen, and oxygen, and their propensity to form [M+NH₄]⁺ adducts contributes to frequent mis-annotation; ceramides and NAE contain a single nitrogen atom and preferentially form [M+H]⁺ adducts. As a result, these structurally distinct lipids may be incorrectly classified with high annotation confidence, despite limited supporting fragmentation evidence. This likely explains the elevated number of DAGs reported in the MS-DIAL-annotated dataset and underscores the need for integrating class-specific fragmentation patterns, such as the “18-series” for wax esters, into existing MS/MS-based annotation platforms to improve compound specificity.

**Figure 7.**
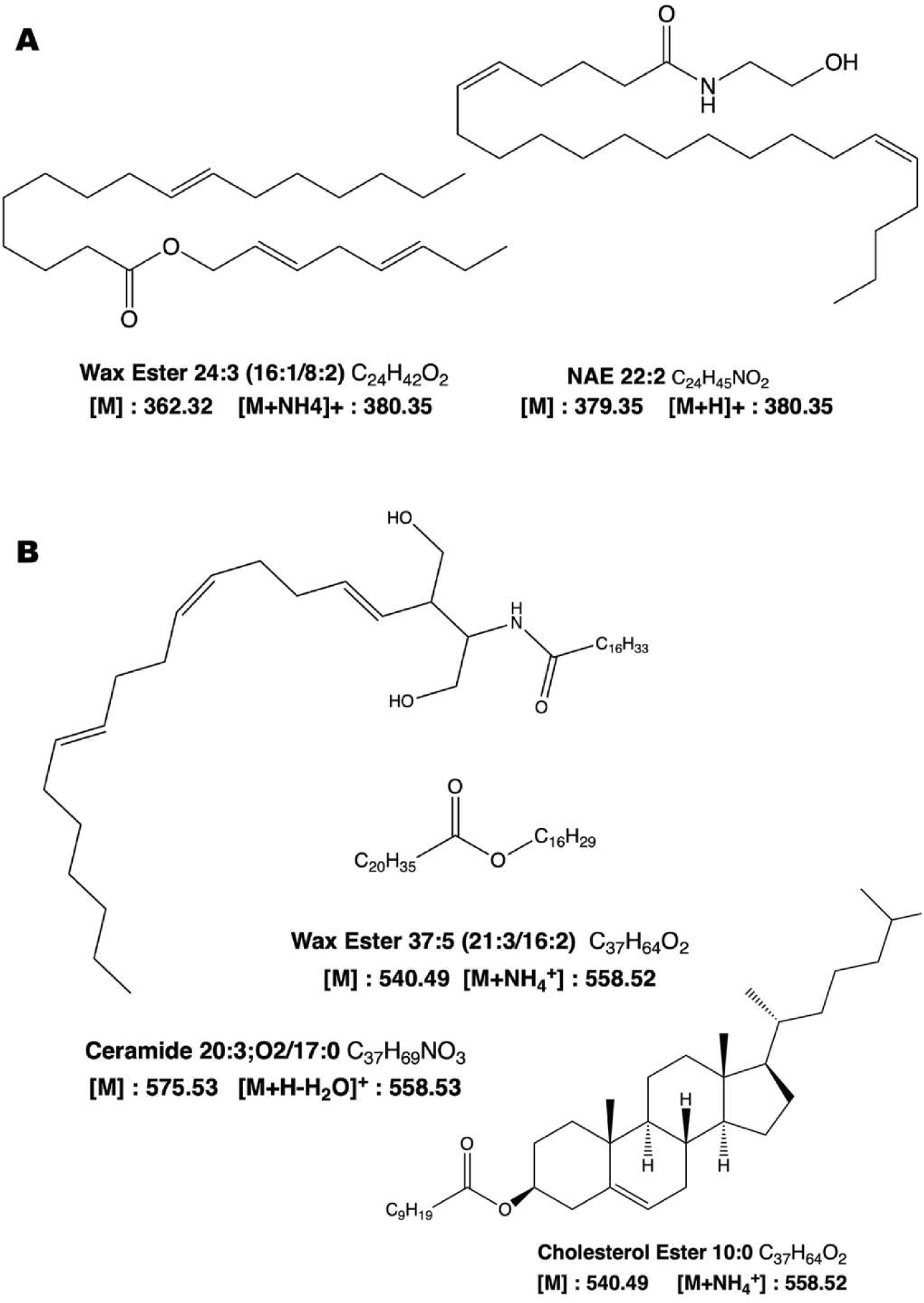
Molecular structures of compounds co-annotated as wax esters. **(A)** Wax ester 24:3 with an ammonium adduct ([M+NH_4_]^+^) and N-Acylethanolamine (NAE) 22:2 with a proton adduct ([M+H]^+^) have the same exact mass (380.35 da) (B) Wax ester 37:5 with an ammonium adduct shares the same mass as Ceramide 20:3+2O with a dehydrated proton adduct ([M+H-H_2_O]^+^) and Cholesterol Ester 10:0 with an ammonium adduct.

## Conclusions

We have presented a robust method for extracting lipids from individuals of trophically important chitin-bound meso-zooplankton, in which lipid metabolism plays a central role. Both meso-zooplankton and their associated lipids are critical not only to marine ecosystems but also to the global carbon cycle. Notably, Jónasdóttir et al. (Jónasdóttir et al., 2015) demonstrated that the seasonal lipid pump, the transport of carbon to the deep ocean via copepod lipids, rivals the particulate sinking flux at high latitudes. In that paper, they use measurements of copepod prosome length and their relationship to oil sac volume to estimate storage lipid content of *C. finmarchicus.* A reliable extraction and analysis protocol for intact copepod lipids would allow for a more detailed understanding of the molecular mechanisms driving this carbon sequestration process.

In particular, our approach enables direct detection of whole molecular species, offering compound-level insight into biological processes. Here, we show that the analysis of the intact lipidomes of individual copepods is not only achievable but also can be easily integrated into pre-existing analysis pipelines, opening a new dimension in this area of research.

Importantly, the efficacy of this specific Bligh and Dyer variation has been well established in phytoplankton and microbial lipidomics (Aveiro et al., 2020; Becker et al., 2018; Bowman et al., 2021; Carriot et al., 2021; Diaz et al., 2021; Laber et al., 2018; Muratore et al., 2022; Popendorf et al., 2013; Rempfert et al., 2023). Expanding its application to mesozooplankton and higher trophic levels enables methodological integration for tracing trophodynamic interactions within marine ecosystems. For example, it may allow us to trace the movement of specific fatty acids from storage lipids in phytoplankton, such as triacylglycerols (TAGs), into zooplankton biomass, including membrane lipids, and other functional lipid classes.

One current approach in zooplankton feeding studies involves gut pigment content analyses, using fluorometric measurements of chlorophyll a and pheophytin to estimate ingestion rates of phytoplankton (Cawley et al., 2021). While fluorometry is straightforward, it is potentially constrained in chemical breadth when used by itself. The addition of intact lipidomics could help quantify the full suite of ingested pigments, such as fucoxanthin, violaxanthin, astaxanthin which allow us to infer phytoplankton community composition of gut content, via tools such as CHEMTAX or *phytoclass* (Hayward et al., 2023; Mackey et al., 1996).

A major advantage of our method is its suitability for capturing biological heterogeneity. Populations at this trophic level and size scale exhibit high inter-individual heterogeneity due to active feeding and mobility (Seuront and Strutton, 2004). We demonstrate that our single-animal extractions capture this variability. In environmental surveys, where individual variability complicates source attribution, samples can be pooled to reduce standard deviations. Conversely, in controlled culture experiments with biological triplicates, single-copepod analyses preserve individual responses to experimental treatments, enabling more nuanced physiological insights.

Finally, from an analytical standpoint, we have identified key limitations in existing lipid annotation pipelines and implemented improvements to better resolve metabolically important compounds, wax esters. Our application of this protocol to both individual and pooled samples enables analyses across multiple biological scales, from capturing the individual-level variability in lab-based experiments to population- or community-scale patterns in environmental samples. This scalability also enables integrative studies across trophic levels, linking lipid metabolism to broader ecosystem functions. Ultimately, our high-resolution, compound-specific approach offers a powerful tool for elucidating the metabolic responses of chitin-bound organisms to environmental change.

---

**Supplementary Figure 1.**
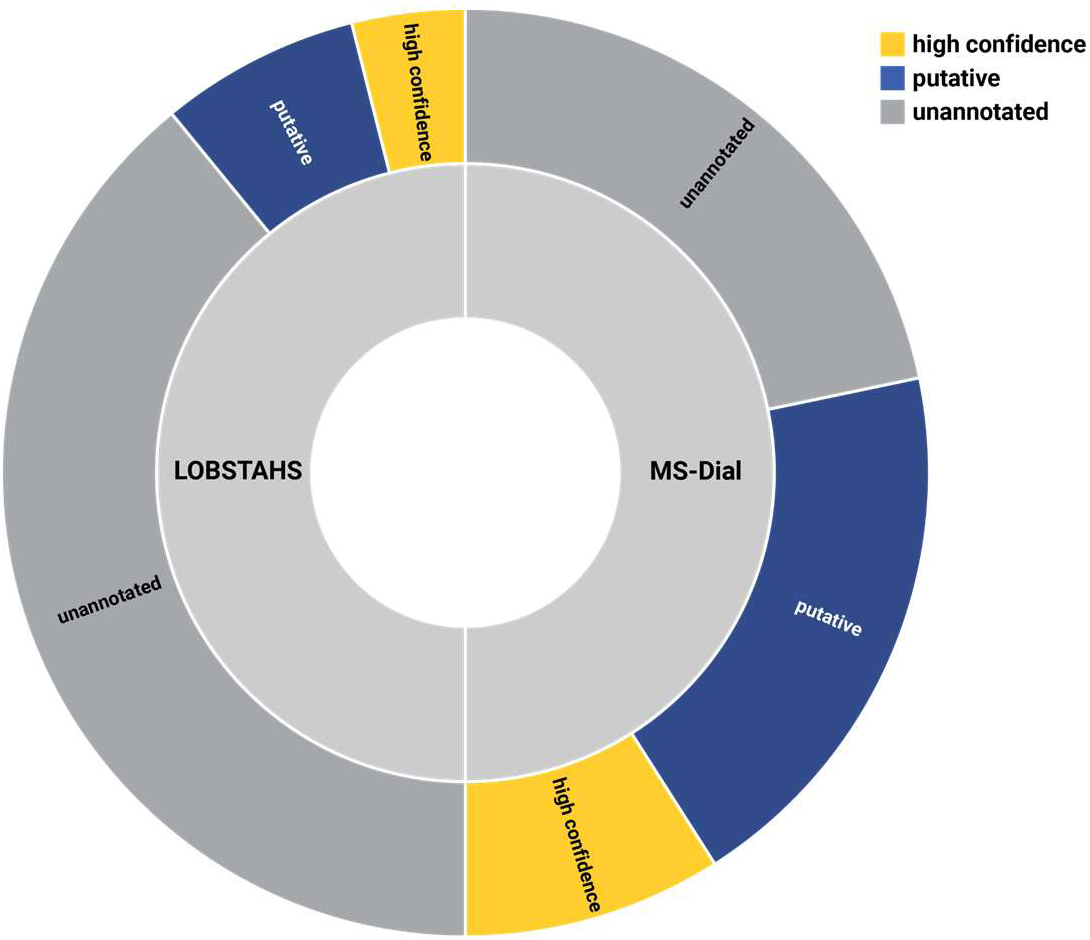
Donut chart of the annotated lipidome based on LOBSTAHS and MS-DIAL. Size of the slices are the number of compounds in each category.

**Supplementary Figure 2.**
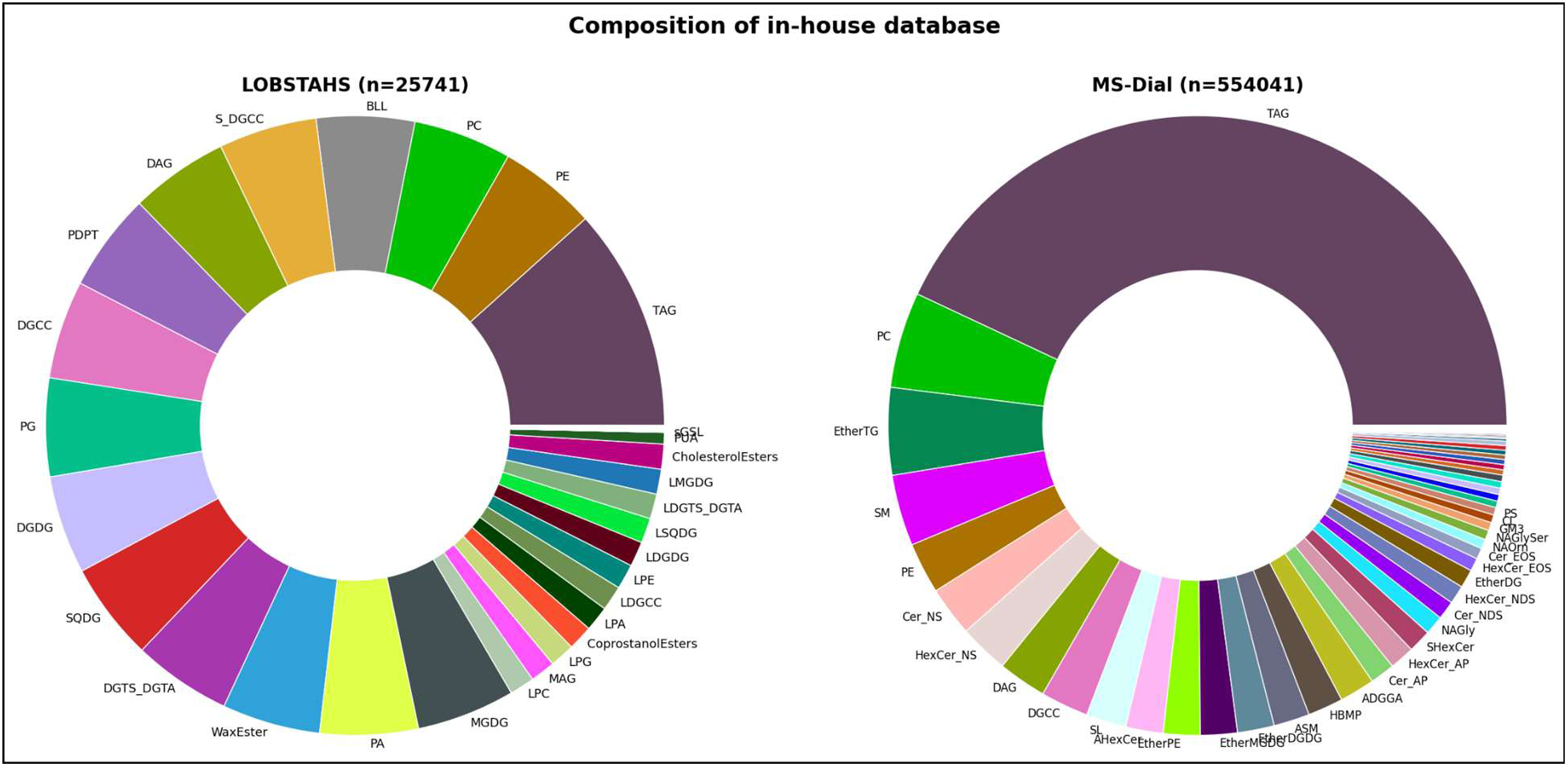
Lipid species composition of the default lipidomic databases in LOBSTAHS and MS-DIAL.

**Supplementary Table 1.**
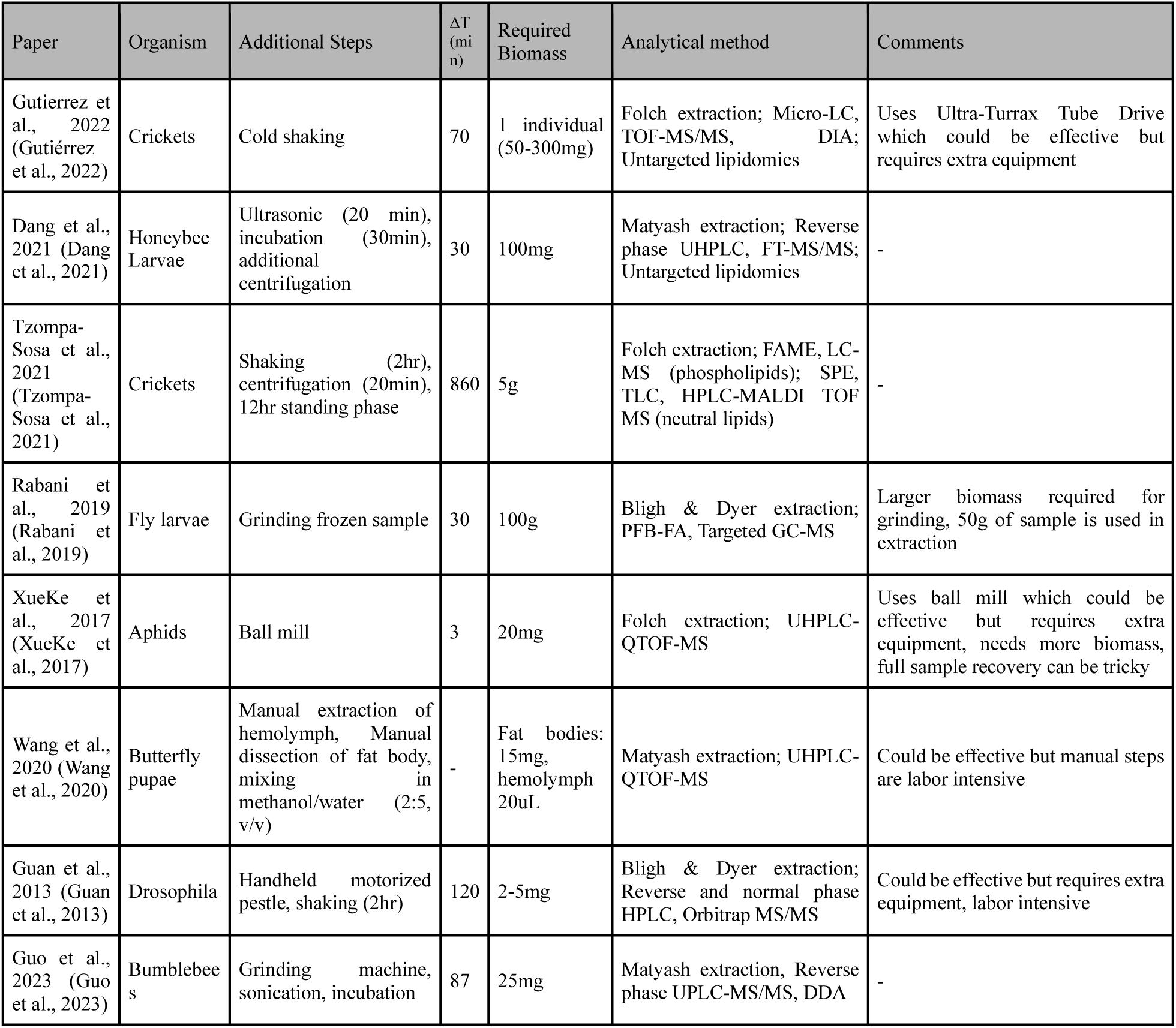
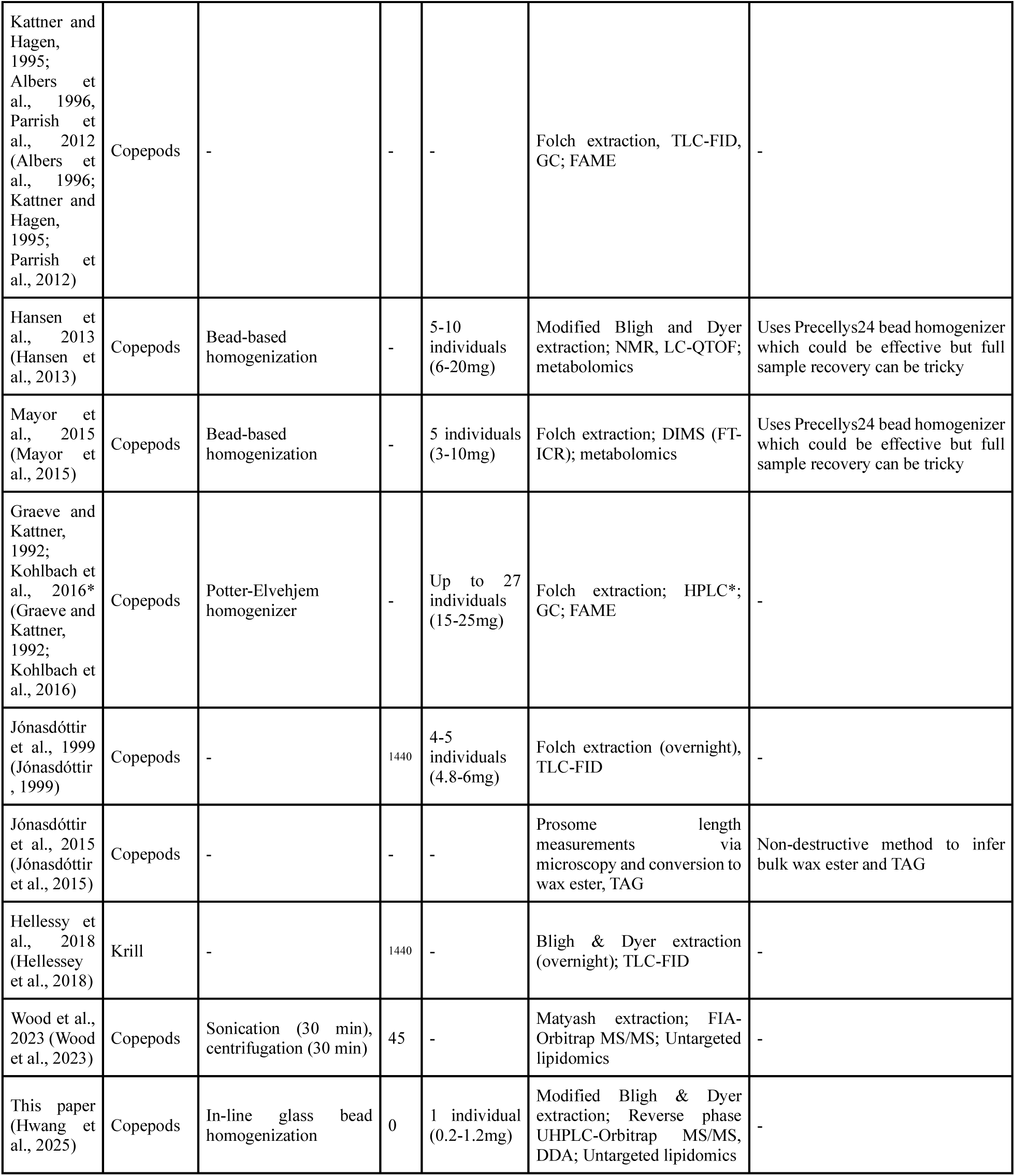
Comparison of lipid extraction methods for chitin-bound organisms. ΔT indicates additional time needed to optimize extraction compared to the method presented in this study. If otherwise not stated in reference text, ΔT and required biomass were denoted with a dash. Extraction method, instrument method, and analytical objective are separated by semicolons when possible. Individual cricket biomass calculations are based on values from Booth and Kiddell (2007) and Lundy and Parrella (2015) (Booth and Kiddell, 2007; Lundy and Parrella, 2015). Individual copepod biomass estimates are based on Aarflot et al. (2018) (Aarflot et al., 2018). Method Abbreviations: DDA – Data-Dependent Acquisition; DIA – Data-Independent Acquisition; DIMS – Direct Infusion Mass Spectrometry; FAME – Fatty Acid Methyl Esters; FIA – Flow Infusion Analysis; FID – Flame Ionization Detection; FT-ICR – Fourier Transform-Ion Cyclotron Resonance; GC – Gas Chromatography; HRMS – High-Resolution Mass Spectrometry; LC – Liquid Chromatography; MALDI – Matrix-Assisted Laser Desorption/Ionization; MS/MS – Tandem Mass Spectrometry; NMR – Nuclear Magnetic Resonance; PFB-FA – Pentafluorobenzyl Fatty Acid; (Q)TOF – Quadrupole Time-of-Flight; SPE – Solid Phase Extraction; TLC – Thin Layer Chromatography; UHPLC – Ultra-High Performance Liquid Chromatography.

**Supplementary Table 2.** Overview of wax ester standards used in optimization of UHPLC-MS analytic method.

| Compound | R'-R<br>(fatty<br>alcohol<br>fatty acid) | Formula | Exact<br>mass |
| --- | --- | --- | --- |
| Wax Ester 36:1 | 20:0-16:1 | C36H70O2 | 534.538 |
| Wax Ester 38:3 | 20:0-18:3 | C38H70O2 | 558.538 |
| Wax Ester 40:4 | 20:0-20:4 | C40H72O2 | 584.553 |
| Wax Ester 38:1 | 22:0-16:1 | C38H74O2 | 562.569 |
| WaxEster 34:1 | 18:1-16:0 | C34H66O1 | 506.506 |
| WaxEster 34:1 | 16:1-18:0 | C34H66O2 | 506.506 |
| WaxEster 34:1 | 18:0-16:1 | C34H66O2 | 506.506 |
| WaxEster 38:1 | 20:0-18:1 | C38H74O2 | 562.569 |
| WaxEster 38:2 | 20:0-18:2 | C38H72O2 | 560.553 |

**Supplementary Table 3.** Comparison of computational load of each peak-picking program. Each program was run on the same dataset (specifically, this paper’s dataset). We monitored process resource use on a Lenovo ThinkPad L13 Yoga Gen 3 laptop running Microsoft Windows 11 Pro (Version 10.0.26100, 64-bit). The system was equipped with a 12th Gen Intel® Core™ i5-1245U processor (10 cores, 12 threads, base frequency 1.60 GHz) and 32 GB of installed RAM. Data and software were run from the primary solid-state drive. Resource logging was performed with a custom PowerShell script that sampled active processes at fixed intervals (every 1s) recording CPU usage, memory allocation, thread count, and runtime. We tracked the executables corresponding to MZmine, MSDIAL, and R processes. Within R, we distinguished between command-line Rscript.exe runs (non-interactive batch execution), the Rsession.exe backend launched by RStudio (interactive IDE engine; UTF-8 variant on Windows), and the Rstudio.exe interface process (IDE shell). For each program, we summarized wall-clock runtime, average CPU utilization, CPU time relative to wall time, peak and average working set memory (RAM). We also recorded maximum thread and handle counts as indicators of system-level load. To ensure comparability, we reported the maximum values observed across all runs of each program, which captures the upper bounds of resource demand for the workflows tested.

| Tool | Runtime (h) | Avg. CPU (%) [max] | CPU vs Wall (%) [max] | Peak RAM (MB) [max] | Avg. RAM (MB) [max] | Threads (max) | Handles (max) |
| --- | --- | --- | --- | --- | --- | --- | --- |
| MS-DIAL | 0.55 | 3.2 | 35.3 | 7353.2 | 5228.5 | 58 | 1524 |
| LOBSTAHS | 5.32 | 9.3 | 96.5 | 4786.0 | 2734.1 | 8 | 243 |
| MZmine | 0.14 | 11.3 | 107.8 | 10809.9 | 3728.7 | 88 | 1690 |

## References

Aarflot, J.M., Skjoldal, H.R., Dalpadado, P., Skern-Mauritzen, M., 2018. Contribution of Calanus species to the mesozooplankton biomass in the Barents Sea. ICES J. Mar. Sci. 75, 2342–2354. 10.1093/icesjms/fsx221

Albers, C.S., Kattner, G., Hagen, W., 1996. The compositions of wax esters, triacylglycerols and phospholipids in Arctic and Antarctic copepods: evidence of energetic adaptations. Mar. Chem. 55, 347–358. 10.1016/S0304-4203(96)00059-X

Aveiro, S.S., Melo, T., Figueiredo, A., Domingues, P., Pereira, H., Maia, I.B., Silva, J., Domingues, M.R., Nunes, C., Moreira, A.S.P., 2020. The Polar Lipidome of Cultured Emiliania huxleyi: A Source of Bioactive Lipids with Relevance for Biotechnological Applications. Biomolecules 10, 1434. 10.3390/biom10101434

Becker, K.W., Collins, J.R., Durham, B.P., Groussman, R.D., White, A.E., Fredricks, H.F., Ossolinski, J.E., Repeta, D.J., Carini, P., Armbrust, E.V., Van Mooy, B.A.S., 2018. Daily changes in phytoplankton lipidomes reveal mechanisms of energy storage in the open ocean. Nat. Commun. 9. 10.1038/s41467-018-07346-z

Booth, D.T., Kiddell, K., 2007. Temperature and the energetics of development in the house cricket (*Acheta domesticus*). J. Insect Physiol. 53, 950–953. 10.1016/j.jinsphys.2007.03.009

Bowman, J.S., Van Mooy, B.A.S., Lowenstein, D.P., Fredricks, H.F., Hansel, C.M., Gast, R., Collins, J.R., Couto, N., Ducklow, H.W., 2021. Whole Community Metatranscriptomes and Lipidomes Reveal Diverse Responses Among Antarctic Phytoplankton to Changing Ice Conditions. Front. Mar. Sci. 8. 10.3389/fmars.2021.593566

Carriot, N., Paix, B., Greff, S., Viguier, B., Briand, J.-F., Culioli, G., 2021. Integration of LC/MS-based molecular networking and classical phytochemical approach allows in-depth annotation of the metabolome of non-model organisms - The case study of the brown seaweed *Taonia atomaria*. Talanta 225, 121925. 10.1016/j.talanta.2020.121925

Cawley, G.F., Décima, M., Mast, A., Prairie, J.C., 2021. The effect of phytoplankton properties on the ingestion of marine snow by *Calanus pacificus*. J. Plankton Res. 43, 957–973. 10.1093/plankt/fbab074

Cequier-Sánchez, E., Rodríguez, C., Ravelo, Á.G., Zárate, R., 2008. Dichloromethane as a Solvent for Lipid Extraction and Assessment of Lipid Classes and Fatty Acids from Samples of Different Natures. J. Agric. Food Chem. 56, 4297–4303. 10.1021/jf073471e

Chen, J., Green, K.B., Nichols, K.K., 2015. Characterization of Wax Esters by Electrospray Ionization Tandem Mass Spectrometry: Double Bond Effect and Unusual Product Ions. Lipids 50, 821–836. 10.1007/s11745-015-4044-6

Dang, X., Li, Y., Li, X., Wang, C., Ma, Z., Wang, L., Fan, X., Li, Z., Huang, D., Xu, J., Zhou, Z., 2021. Lipidomic Profiling Reveals Distinct Differences in Sphingolipids Metabolic Pathway between Healthy Apis cerana cerana larvae and Chinese Sacbrood Disease. Insects 12, 703. 10.3390/insects12080703

Diaz, B.P., Knowles, B., Johns, C.T., Laber, C.P., Bondoc, K.G.V., Haramaty, L., Natale, F., Harvey, E.L., Kramer, S.J., Bolaños, L.M., Lowenstein, D.P., Fredricks, H.F., Graff, J., Westberry, T.K., Mojica, K.D.A., Haëntjens, N., Baetge, N., Gaube, P., Boss, E., Carlson, C.A., Behrenfeld, M.J., Van Mooy, B.A.S., Bidle, K.D., 2021. Seasonal mixed layer depth shapes phytoplankton physiology, viral production, and accumulation in the North Atlantic. Nat. Commun. 12, 6634. 10.1038/s41467-021-26836-1

Graeve, M., Kattner, G., 1992. Species-specific differences in intact wax esters of *Calanus hyperboreus* and *C. finmarchicus* from Fram Strait — Greenland Sea. Mar. Chem. 39, 269–281. 10.1016/0304-4203(92)90013-Z

Guan, X.L., Cestra, G., Shui, G., Kuhrs, A., Schittenhelm, R.B., Hafen, E., van der Goot, F.G., Robinett, C.C., Gatti, M., Gonzalez-Gaitan, M., Wenk, M.R., 2013. Biochemical Membrane Lipidomics during Drosophila Development. Dev. Cell 24, 98–111. 10.1016/j.devcel.2012.11.012

Guo, Yueqin, Liu, F., Guo, Yulong, Qu, Y., Zhang, Z., Yao, J., Xu, J., Li, J., 2023. Untargeted Lipidomics Analysis Unravels the Different Metabolites in the Fat Body of Mated Bumblebee (Bombus terrestris) Queens. Int. J. Mol. Sci. 24, 15408. 10.3390/ijms242015408

Gutiérrez, Y., Fresch, M., Scherber, C., Brockmeyer, J., 2022. The lipidome of an omnivorous insect responds to diet composition and social environment. Ecol. Evol. 12, e9497. 10.1002/ece3.9497

Hansen, B.H., Degnes, K., Øverjordet, I.B., Altin, D., Størseth, T.R., 2013. Metabolic fingerprinting of arctic copepods Calanus finmarchicus, Calanus glacialis and Calanus hyperboreus. Polar Biol. 36, 1577–1586. 10.1007/s00300-013-1375-8

Hayward, A., Pinkerton, M.H., Gutierrez-Rodriguez, A., 2023. phytoclass: A pigment-based chemotaxonomic method to determine the biomass of phytoplankton classes. Limnol. Oceanogr. Methods 21, 220–241. 10.1002/lom3.10541

Hellessey, N., Ericson, J.A., Nichols, P.D., Kawaguchi, S., Nicol, S., Hoem, N., Virtue, P., 2018. Seasonal and interannual variation in the lipid content and composition of Euphausia superba Dana, 1850 (Euphausiacea) samples derived from the Scotia Sea fishery. J. Crustac. Biol. 38, 673–681. 10.1093/jcbiol/ruy053

Hummel, J., Segu, S., Li, Y., Irgang, S., Jueppner, J., Giavalisco, P., 2011. Ultra performance liquid chromatography and high resolution mass spectrometry for the analysis of plant lipids. Front. Plant Sci. 2, 1–17. 10.3389/fpls.2011.00054

Hunter, J.E., Fredricks, H.F., Behrendt, L., Alcolombri, U., Bent, S.M., Stocker, R., Van Mooy, B.A.S., 2021. Using High-Sensitivity Lipidomics To Assess Microscale Heterogeneity in Oceanic Sinking Particles and Single Phytoplankton Cells. Environ. Sci. Technol. 55, 15456–15465. 10.1021/acs.est.1c02836

Hwang, J., Hayward, A., Sofen, L.E., Pitz, K.J., Chavez, F.P., Edwards, B.R., 2025. Daily microbial rhythms of the surface ocean interrupted by the new moon—a lipidomic study. ISME Commun. 5, ycaf044. 10.1093/ismeco/ycaf044

Jónasdóttir, S.H., 1999. Lipid content of Calanus finmarchicus during overwintering in the Faroe-Shetland Channel. Fish. Oceanogr. 8, 61–72. 10.1046/j.1365-2419.1999.00003.x

Jónasdóttir, S.H., Visser, A.W., Richardson, K., Heath, M.R., 2015. Seasonal copepod lipid pump promotes carbon sequestration in the deep North Atlantic. Proc. Natl. Acad. Sci. 112, 12122–12126. 10.1073/pnas.1512110112

Kattner, G., Hagen, W., 1995. Polar herbivorous copepods - different pathways in lipid biosynthesis. ICES J. Mar. Sci. 52, 329–335. 10.1016/1054-3139(95)80048-4

Kind, T., Liu, K.-H., Lee, D.Y., DeFelice, B., Meissen, J.K., Fiehn, O., 2013. LipidBlast in silico tandem mass spectrometry database for lipid identification. Nat. Methods 10, 755–758. 10.1038/nmeth.2551

Kiyonami, R., Peake, David A, Yokoi, Yasuto, Miller, Ken, n.d. Increased Throughput and Confidence for Lipidomics Profiling Using Comprehensive HCD MS2 and CID MS2/MS3 on a Tribrid Orbitrap Mass Spectrometer 10.

Kohlbach, D., Graeve, M., A. Lange, B., David, C., Peeken, I., Flores, H., 2016. The importance of ice algae-produced carbon in the central Arctic Ocean ecosystem: Food web relationships revealed by lipid and stable isotope analyses. Limnol. Oceanogr. 61, 2027–2044. 10.1002/lno.10351

Laber, C.P., Hunter, J.E., Carvalho, F., Collins, J.R., Hunter, E.J., Schieler, B.M., Boss, E., More, K., Frada, M., Thamatrakoln, K., Brown, C.M., Haramaty, L., Ossolinski, J., Fredricks, H., Nissimov, J.I., Vandzura, R., Sheyn, U., Lehahn, Y., Chant, R.J., Martins, A.M., Coolen, M.J.L., Vardi, A., Ditullio, G.R., Van Mooy, B.A.S., Bidle, K.D., 2018. Coccolithovirus facilitation of carbon export in the North Atlantic. Nat. Microbiol. 3, 537–547. 10.1038/s41564-018-0128-4

Lee, J., Tantillo, D.J., Wang, L.-P., Fiehn, O., 2024. Predicting Collision-Induced-Dissociation Tandem Mass Spectra (CID-MS/MS) Using Ab Initio Molecular Dynamics. J. Chem. Inf. Model. 64, 7470–7487. 10.1021/acs.jcim.4c00760

Lundy, M.E., Parrella, M.P., 2015. Crickets Are Not a Free Lunch: Protein Capture from Scalable Organic Side-Streams via High-Density Populations of Acheta domesticus. PLOS ONE 10, e0118785. 10.1371/journal.pone.0118785

Mackey, M.D., Mackey, D.J., Higgins, H.W., Wright, S.W., 1996. CHEMTAX - A program for estimating class abundances from chemical markers: Application to HPLC measurements of phytoplankton. Mar. Ecol. Prog. Ser. 144, 265–283. 10.3354/meps144265

Matyash, V., Liebisch, G., Kurzchalia, T.V., Shevchenko, A., Schwudke, D., 2008. Lipid extraction by methyl-tert-butyl ether for high-throughput lipidomics. J. Lipid Res. 49, 1137–1146. 10.1194/jlr.D700041-JLR200

Mayor, D.J., Sommer, U., Cook, K.B., Viant, M.R., 2015. The metabolic response of marine copepods to environmental warming and ocean acidification in the absence of food. Sci. Rep. 5, 13690. 10.1038/srep13690

Muratore, D., Boysen, A.K., Harke, M.J., Becker, K.W., Casey, J.R., Coesel, S.N., Mende, D.R., Wilson, S.T., Aylward, F.O., Eppley, J.M., Vislova, A., Peng, S., Rodriguez-Gonzalez, R.A., Beckett, S.J., Virginia Armbrust, E., DeLong, E.F., Karl, D.M., White, A.E., Zehr, J.P., Van Mooy, B.A.S., Dyhrman, S.T., Ingalls, A.E., Weitz, J.S., 2022. Complex marine microbial communities partition metabolism of scarce resources over the diel cycle. Nat. Ecol. Evol. 6, 218–229. 10.1038/s41559-021-01606-w

Murtagh, F., Legendre, P., 2014. Ward’s Hierarchical Agglomerative Clustering Method: Which Algorithms Implement Ward’s Criterion? J. Classif. 31, 274–295. 10.1007/s00357-014-9161-z

Parrish, C.C., French, V.M., Whiticar, M.J., 2012. Lipid class and fatty acid composition of copepods (Calanus finmarchicus, C. glacialis, Pseudocalanus sp., Tisbe furcata and Nitokra lacustris) fed various combinations of autotrophic and heterotrophic protists. J. Plankton Res. 34, 356–375. 10.1093/plankt/fbs003

Popendorf, K.J., Fredricks, H.F., Van Mooy, B.A.S., 2013. Molecular Ion-Independent Quantification of Polar Glycerolipid Classes in Marine Plankton Using Triple Quadrupole MS. Lipids 48, 185–195. 10.1007/s11745-012-3748-0

Rabani, V., Cheatsazan, H., Davani, S., 2019. Proteomics and Lipidomics of Black Soldier Fly (Diptera: Stratiomyidae) and Blow Fly (Diptera: Calliphoridae) Larvae. J. Insect Sci. 19, 29. 10.1093/jisesa/iez050

Rempfert, K.R., Kraus, E.A., Nothaft, D.B., Dildar, N., Spear, J.R., Sepúlveda, J., Templeton, A.S., 2023. Intact polar lipidome and membrane adaptations of microbial communities inhabiting serpentinite-hosted fluids. Front. Microbiol. 14. 10.3389/fmicb.2023.1198786

Rinaudo, M., 2006. Chitin and chitosan: Properties and applications. Prog. Polym. Sci. 31, 603–632. 10.1016/j.progpolymsci.2006.06.001

Sargent, J.R., Gatten, R.R., McIntosh, R., 1977. Wax esters in the marine environment - their occurrence, formation, transformation and ultimate fates. Mar. Chem. 5, 573–584. 10.1016/0304-4203(77)90043-3

Schmid, R., Heuckeroth, S., Korf, A., Smirnov, A., Myers, O., Dyrlund, T.S., Bushuiev, R., Murray, K.J., Hoffmann, N., Lu, M., Sarvepalli, A., Zhang, Z., Fleischauer, M., Dührkop, K., Wesner, M., Hoogstra, S.J., Rudt, E., Mokshyna, O., Brungs, C., Ponomarov, K., Mutabdžija, L., Damiani, T., Pudney, C.J., Earll, M., Helmer, P.O., Fallon, T.R., Schulze, T., Rivas-Ubach, A., Bilbao, A., Richter, H., Nothias, L.-F., Wang, M., Orešič, M., Weng, J.-K., Böcker, S., Jeibmann, A., Hayen, H., Karst, U., Dorrestein, P.C., Petras, D., Du, X., Pluskal, T., 2023. Integrative analysis of multimodal mass spectrometry data in MZmine 3. Nat. Biotechnol. 41, 447–449. 10.1038/s41587-023-01690-2

Seuront, L., Strutton, P.G., 2004. Using Multiagent Systems to Develop Individual-Based Models for Copepods: Consequences of Individual Behavior and Spatial Heterogeneity on the Emerging Properties at the Population Scale, in: Handbook of Scaling Methods in Aquatic Ecology. CRC Press, United Kingdom, pp. 543–562. 10.1201/9780203489550-42

Sturt, H.F., Summons, R.E., Smith, K., Elvert, M., Hinrichs, K.U., 2004. Intact polar membrane lipids in prokaryotes and sediments deciphered by high-performance liquid chromatography/electrospray ionization multistage mass spectrometry - New biomarkers for biogeochemistry and microbial ecology. Rapid Commun. Mass Spectrom. 18, 617–628. 10.1002/rcm.1378

Sumner, L.W., Amberg, A., Barrett, D., Beale, M.H., Beger, R., Daykin, C.A., Fan, T.W.-M., Fiehn, O., Goodacre, R., Griffin, J.L., Hankemeier, T., Hardy, N., Harnly, J., Higashi, R., Kopka, J., Lane, A.N., Lindon, J.C., Marriott, P., Nicholls, A.W., Reily, M.D., Thaden, J.J., Viant, M.R., 2007. Proposed minimum reporting standards for chemical analysis Chemical Analysis Working Group (CAWG) Metabolomics Standards Initiative (MSI). Metabolomics Off. J. Metabolomic Soc. 3, 211–221. 10.1007/s11306-007-0082-2

Tsugawa, H., Ikeda, K., Takahashi, M., Satoh, A., Mori, Y., Uchino, H., Okahashi, N., Yamada, Y., Tada, I., Bonini, P., Higashi, Y., Okazaki, Y., Zhou, Z., Zhu, Z.J., Koelmel, J., Cajka, T., Fiehn, O., Saito, K., Arita, Masanori, Arita, Makoto, 2020. A lipidome atlas in MS-DIAL 4. Nat. Biotechnol. 38, 1159–1163. 10.1038/s41587-020-0531-2

Tsugawa, H., Ikeda, K., Tanaka, W., Senoo, Y., Arita, Makoto, Arita, Masanori, 2017. Comprehensive identification of sphingolipid species by in silico retention time and tandem mass spectral library. J. Cheminformatics 9, 19. 10.1186/s13321-017-0205-3

Tzompa-Sosa, D.A., Dewettinck, K., Provijn, P., Brouwers, J.F., De Meulenaer, B., Oonincx, D.G.A.B., 2021. Lipidome of cricket species used as food. Food Chem. 349, 129077. 10.1016/j.foodchem.2021.129077

Van Mooy, B. a. S., Moutin, T., Duhamel, S., Rimmelin, P., Van Wambeke, F., 2008. Phospholipid synthesis rates in the eastern subtropical South Pacific Ocean. Biogeosciences 5, 133–139. 10.5194/bg-5-133-2008

Van Mooy, B.A.S., Fredricks, H.F., 2010. Bacterial and eukaryotic intact polar lipids in the eastern subtropical South Pacific: Water-column distribution, planktonic sources, and fatty acid composition. Geochim. Cosmochim. Acta 74, 6499–6516. 10.1016/j.gca.2010.08.026

Wang, J., Jin, H., Schlenke, T., Yang, Y., Wang, F., Yao, H., Fang, Q., Ye, G., 2020. Lipidomics reveals how the endoparasitoid wasp *Pteromalus puparum* manipulates host energy stores for its young. Biochim. Biophys. Acta BBA - Mol. Cell Biol. Lipids 1865, 158736. 10.1016/j.bbalip.2020.158736

Wang, M., Carver, J.J., Phelan, V.V., Sanchez, L.M., Garg, N., Peng, Y., Nguyen, D.D., Watrous, J., Kapono, C.A., Luzzatto-Knaan, T., Porto, C., Bouslimani, A., Melnik, A.V., Meehan, M.J., Liu, W.-T., Crüsemann, M., Boudreau, P.D., Esquenazi, E., Sandoval-Calderón, M., Kersten, R.D., Pace, L.A., Quinn, R.A., Duncan, K.R., Hsu, C.-C., Floros, D.J., Gavilan, R.G., Kleigrewe, K., Northen, T., Dutton, R.J., Parrot, D., Carlson, E.E., Aigle, B., Michelsen, C.F., Jelsbak, L., Sohlenkamp, C., Pevzner, P., Edlund, A., McLean, J., Piel, J., Murphy, B.T., Gerwick, L., Liaw, C.-C., Yang, Y.-L., Humpf, H.-U., Maansson, M., Keyzers, R.A., Sims, A.C., Johnson, A.R., Sidebottom, A.M., Sedio, B.E., Klitgaard, A., Larson, C.B., Boya P, C.A., Torres-Mendoza, D., Gonzalez, D.J., Silva, D.B., Marques, L.M., Demarque, D.P., Pociute, E., O’Neill, E.C., Briand, E., Helfrich, E.J.N., Granatosky, E.A., Glukhov, E., Ryffel, F., Houson, H., Mohimani, H., Kharbush, J.J., Zeng, Y., Vorholt, J.A., Kurita, K.L., Charusanti, P., McPhail, K.L., Nielsen, K.F., Vuong, L., Elfeki, M., Traxler, M.F., Engene, N., Koyama, N., Vining, O.B., Baric, R., Silva, R.R., Mascuch, S.J., Tomasi, S., Jenkins, S., Macherla, V., Hoffman, T., Agarwal, V., Williams, P.G., Dai, J., Neupane, R., Gurr, J., Rodríguez, A.M.C., Lamsa, A., Zhang, C., Dorrestein, K., Duggan, B.M., Almaliti, J., Allard, P.-M., Phapale, P., Nothias, L.-F., Alexandrov, T., Litaudon, M., Wolfender, J.-L., Kyle, J.E., Metz, T.O., Peryea, T., Nguyen, D.-T., VanLeer, D., Shinn, P., Jadhav, A., Müller, R., Waters, K.M., Shi, W., Liu, X., Zhang, L., Knight, R., Jensen, P.R., Palsson, B.Ø., Pogliano, K., Linington, R.G., Gutiérrez, M., Lopes, N.P., Gerwick, W.H., Moore, B.S., Dorrestein, P.C., Bandeira, N., 2016. Sharing and community curation of mass spectrometry data with Global Natural Products Social Molecular Networking. Nat. Biotechnol. 34, 828–837. 10.1038/nbt.3597

Wang, Z., Cao, M., Lam, S.M., Shui, G., 2023. Embracing lipidomics at single-cell resolution: Promises and pitfalls. TrAC Trends Anal. Chem. 160, 116973. 10.1016/j.trac.2023.116973

Wood, P.L., Wood, M.D., Kunigelis, S.C., 2023. Pilot Lipidomics Study of Copepods: Investigation of Potential Lipid-Based Biomarkers for the Early Detection and Quantification of the Biological Effects of Climate Change on the Oceanic Food Chain. Life 13, 2335. 10.3390/life13122335

XueKe, G., Shuai, Z., JunYu, L., LiMin, L., LiJuan, Z., JinJie, C., 2017. Lipidomics and RNA-Seq Study of Lipid Regulation in Aphis gossypii parasitized by Lysiphlebia japonica. Sci. Rep. 7, 1364. 10.1038/s41598-017-01546-1

